# Retention of an endosymbiont for the production of a single molecule

**DOI:** 10.1101/2024.01.04.574232

**Authors:** Arkadiy I. Garber, Andrés Garcia de la Filia Molina, Isabelle Vea, Andrew J. Mongue, Laura Ross, John P. McCutcheon

## Abstract

Sap-feeding insects often maintain multiple nutritional endosymbionts, which act in concert to produce compounds essential for insect survival. Many mealybugs have endosymbionts in a nested configuration: one or two bacterial species reside within the cytoplasm of another bacterium, and together these bacteria have genomes which encode interdependent but complete sets of genes needed to produce key nutritional molecules. Here we show that the mealybug *Pseudococcus viburni* has three endosymbionts, one of which contributes only two genes that produce a single host nutrition-related molecule. All three bacterial endosymbionts have tiny genomes, suggesting that they have been co-evolving inside their insect host for millions of years.

**Significance:** Nutritional endosymbionts synthesize (or contribute to the synthesis of) key metabolites such as essential amino acids and vitamins for their host organism. These nutrients are required by hosts because of their restricted diets, which in the case of mealybugs consists solely of plant phloem sap. Genome sequencing of insect endosymbionts has shown that their genomes can be very small, encoding few genes outside of core bacterial processes and nutrient provisioning. Here we highlight an example that has taken this reductive process to the extreme: a mealybug endosymbiont contributes only a single essential compound, chorismate, to the symbiosis.

## Introduction

Sap-feeding insects form long-term endosymbioses with bacteria or fungi to supplement their diets with essential amino acids and vitamins (Baumann, 2005). Bacteria that form endosymbioses undergo stereotyped and sometimes extreme genome reduction during co-evolution with their insect hosts (McCutcheon and Moran, 2011). Endosymbionts are sometimes supplemented or replacement by new bacterial or fungal symbionts (Koga and Moran, 2014; Matsuura et al., 2018; Husnik and McCutcheon, 2016; Dial et al., 2021). In mealybugs (Hemiptera: Pseudococcidae), as in other related insects (Bennett and Moran, 2013; Oakeson et al., 2014; Mao and Bennett, 2020), symbiont replacement and supplementation has occurred multiple times, resulting in a diversity of symbiont types and ages across species (Husnik and McCutcheon, 2016).

For example, in the handful of mealybug species with available genomic data, numerous bacterial symbionts in the Sodalis genus have been found whose genomes range in size over an order of magnitude, from 3.7 megabase pairs (Mb) (Garber et al., 2021) to 0.35 Mb (Husnik and McCutcheon, 2016). It is thought that this variation in genome size reflects variation in endosymbiont age: newly established endosymbionts have larger genomes, and endosymbionts that have had long associations have smaller genomes (Moran, 1996; Andereson and Kurland, 1998; Andereson and Anderson, 1998; Wernegreen, 2002; Moran et al., 2009; Wolf and Koonin 2013; Oakeson et al., 2014).

In most sequenced mealybugs, a single *Sodalis* endosymbiont resides within the cytoplasm of another bacterial endosymbiont, *Tremblaya princeps* (von Dohlen et al., 2001). There has been one report of a mealybug with two intra-Tremblaya endosymbionts, both with large genomes and likely recently acquired (Garber et al., 2021). Here, we report a similar three-way endosymbiosis, but where all symbionts have highly reduced genomes and so we infer that they have been co-evolving with their host insect for millions of years. Remarkably, one endosymbiont provides only one unique nutrition-related molecule to the symbiosis.

## Results and Discussion

### Endosymbiont genome assembly

Hybrid assembly of endosymbiont contigs using PacBio and Illumina reads resulted in four circular-mapping contigs, two of which (754,563 bp and 281,389 bp in length) are affiliated with the Sodalis group within Gammaproteobacteria. The other two circular contigs (123,124 bp and 20,943 bp in length) belong to *Tremblaya princeps*. Combined, the two *Tremblaya* contigs add up to the typical size of Tremblaya’s genome (144 Kb) from other mealybug species (Husnik and McCutcheon, 2016). It is unclear how *Tremblaya’s* genome has fragmented into two circles, but genome instability is not uncommon in endosymbionts (Van Leuven et al., 2014; Campbell et al., 2015; Campbell et al., 2017) and mitochondria (Palmer and Shields et al., 1984; Vlcek et al., 2011; Shao et al., 2017; Shao et al., 2012; Wu et al., 2015). Read mapping revealed that both gammaproteobacterial contigs have distinct but similar read coverages (81x and 104x). Tremblaya has a much higher read coverage (1798x), and likely maintains many copies of its genome, as reported in the *Tremblaya* symbiont of the long-tailed mealybug, *Pseudococcus longispinus* (Garber et al., 2021), and in the obligate intracellular symbionts of other insects (Van Leuven et al., 2014, Woyke et al., 2010, Komaki and Ishikawa, 1999).

### Pseudococcus viburni harbors two ancient Sodalis-related endosymbionts

Each Sodalis-related contig encodes its own complete set of ribosomal proteins, tRNA genes, and rRNA genes (**Supplemental Figure 1**). The larger 755 Kb contig encodes two copies of the rRNA operon (**Supplemental Figure 2**). A phylogenomic tree (**Figure 1**) supports the presence of two species of *Sodalis* symbionts, with one endosymbiont (755 Kb) clustering with *Moranella endobia* (hereafter, *Moranella*) (McCutcheon and von Dohlen, 2011), and the other (281 Kb) branching off from the phylogenetic cluster that encompasses *Mikella endobia* (Husnik and Mc-Cutcheon, 2016) and *Trabutinella endobia* (Szabó et al., 2017). The similar read coverage depth of each *Sodalis*-related endosymbiont suggests that cells from both symbiont species are present at similar abundances.

**Figure 1.**
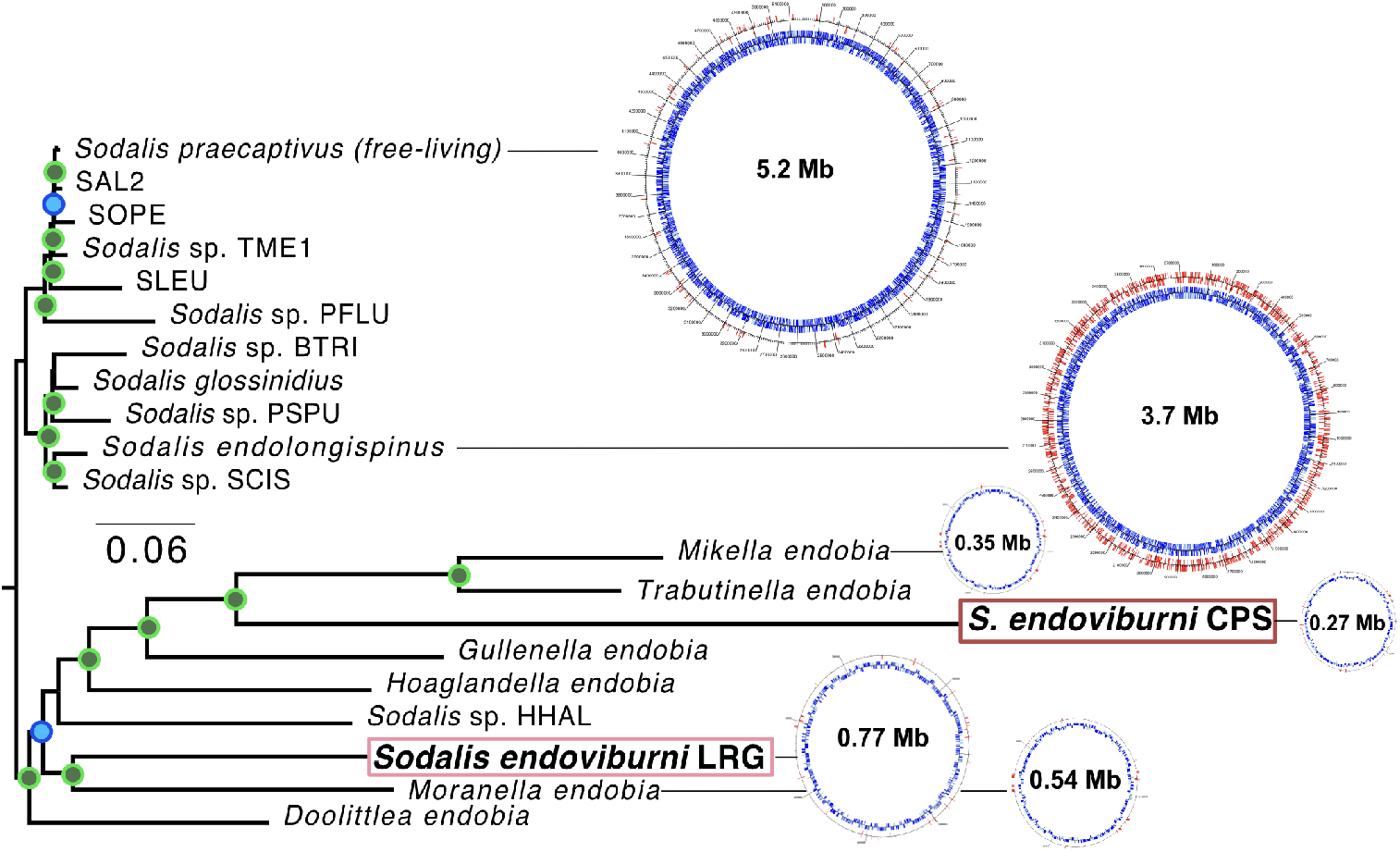
Phylogenomic tree showing the relationship of the two P. viburni gammaproteobacterial endosymbionts (highlighted within dark and light red boxes) with other members from the Sodalis clade. Genome maps from select Sodalis-related endosymbionts, as well as the free-living S. praecaptivus, are shown. Numbers inside each genome map show the size of the genome in mega-bases (million bases); genome maps are divided into two tracks, with the blue inner track showing the locations of protein-coding genes, and the other red track shows the locations of predicted pseudogenes. Nodes with 99% or more support are designated with filled green circles. Nodes with support values between 80% and 98% are colored blue. Nodes with less than 80% support are unlabaled.

**Figure 2.**
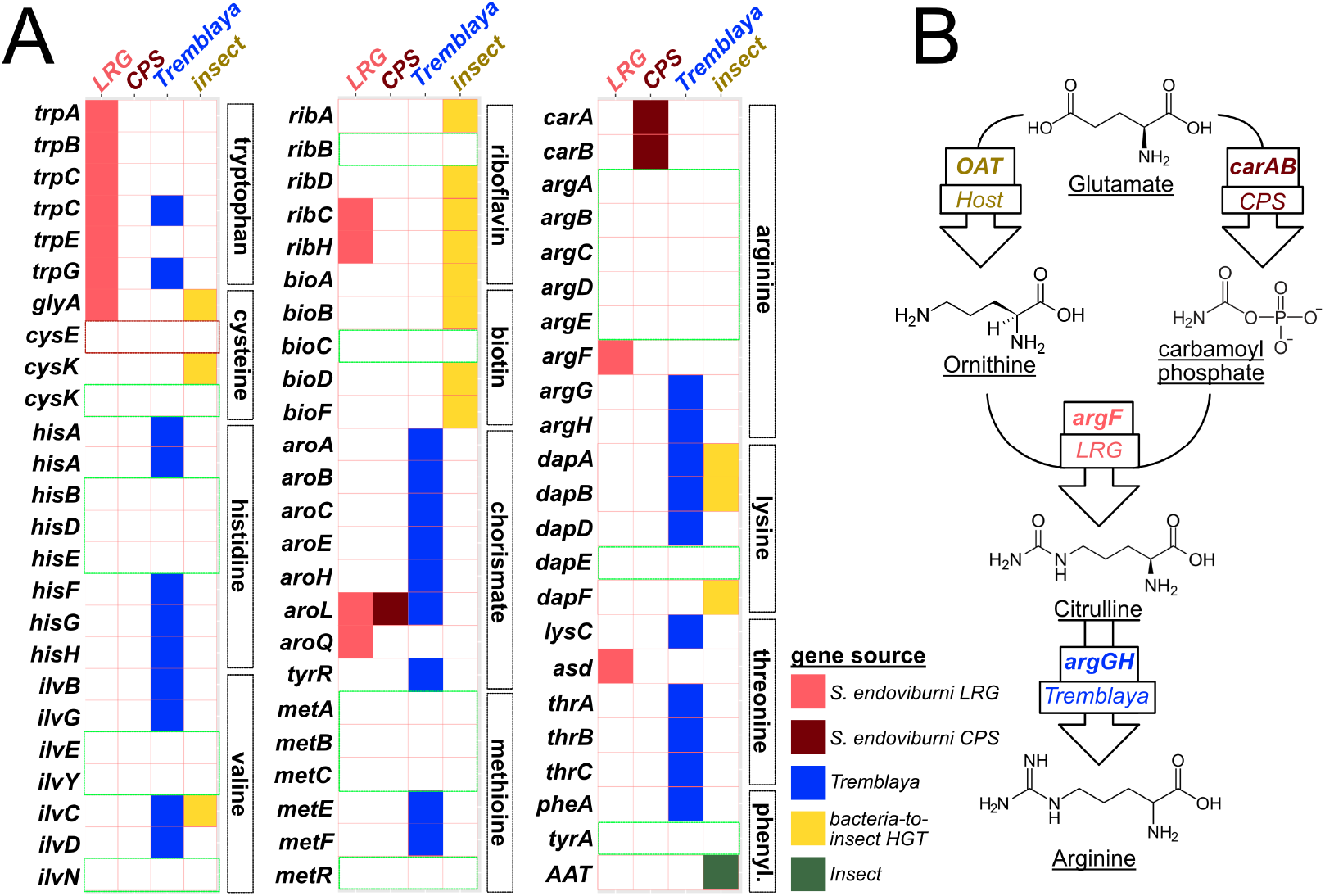
A) presence/absence matrix showing the partitioning of biosynthetic pathway components across the P. viburni symbionts and host. The two Sodalis-related endosymbionts are denoted simply with LRG and CPS. B) Diagram of arginine biosynthesis is P. viburni, showing the sole role of S. endoviburni CPS in generating the carbamoyl phosphate that’s essential of the synthesis of citrulline, a precursor of arginine. Green boxes represent pathway components that are missing in both P. viburni and P. citri mealybugs; red boxes represent pathway components that are missing only in P. viburni.

The two *Sodalis* endosymbionts have highly reduced and gene-dense genomes, with relatively few pseudogenes (<10% of total genes). These features, along with long branch lengths in the phylogenomic tree, suggest that both *Sodalis*-related endosymbionts are ancient (Moran, 1996; Wernegreen, 2002, McCutcheon and Moran, 2011).

### Naming of the novel Sodalis-related symbionts

We propose the name *Candidatus* Sodalis endoviburni LRG (hereafter *S. endovirburni* LRG) for the *Sodalis*-allied organism with the larger genome (LRG meaning large) and *Candidatus* Sodalis endoviburni CPS (hereafter *S. endovirburni* CPS) for the *Sodalis*-allied organism with the smaller genome (CPS reflecting that all this organism seems to contribute to the symbiosis is carbamoyl phosphate synthesis, see the next section for a description of this genome).

### Carbamoyl phosphate synthase: S. endoviburni CPS’s only contribution to the symbiosis

To examine nutritional contributions and metabolic complementarity between the two *Sodalis* endosymbionts of *P. viburni*, we screened both genomes, along with *Tremblaya* and the host’s genome, for pathways relevant to amino acid and vitamin biosynthesis (Bauman, 2005; Douglas, 2006). We found that genes for these pathways are mostly retained on the genomes of *S. endoviburni* LRG, *Tremblaya*, and the host (**Figure 2A**). The nuclear genome of *P. viburni*, like the closely related mealybugs *Pseudococcus longispinus* and *Planococcus citri*, encodes numerous bacterial genes (acquired via horizonal gene transfer) that seem to complement genes missing from the bacterial symbiont genomes (Husnik and McCutcheon, 2016 Bublitz et al., 2019). Our screen identified the same HGTs in *P. viburni* that were previously reported in the citrus mealybug *P. citri* (Husnik et al., 2013), suggesting these HGT events occurred prior to the split between *Pseudococcus* and *Planococcus*. Surprisingly, *S. endoviburni* CPS seems to only contribute three genes related to the biosynthetic pathways for essential amino acids: the small subunit of carbamoyl-phosphate synthase (*carA*), the large subunit of carbamoyl-phosphate synthase (*carB*), and shikimate kinase II (*aroL*). While *aroL* is essential for the synthesis of chorismate and subsequently a number of aromatic amino acids, it is also present in the genomes of *S. endoviburni* LRG and *Tremblaya*. It thus appears that the only unique nutritional contribution from *S. endoviburni* CPS is carbamoyl-phosphate (from *carAB*), used in the production of the essential amino acid arginine (**Figure 2B**).

While *S. endoviburni* CPS represents the smallest genome within the *Sodalis* clade of symbionts, it is not the smallest symbiont genome sequenced so far. Smaller still are the symbionts of some sap-feeding leafhoppers, which have bacterial endosymbionts with genomes as small as about 100 kb, yet still encode more than three genes for the biosynthesis of essential metabolites from the insects’ sugar-based diet (Bennett and Moran, 2013). Two other examples rival the level of specialization we report here for *S. endoviburni* CPS. The first is the ancient symbiont *Stammera* of the plant-feeding leaf beetle, which only encodes a few genes required for the breakdown of pectin (Salem et al., 2017). The second is a case in which an endosymbiont genome appears to retain no symbiotic genes at all, but rather seems to have eroded to the point of being nutritionally useless and likely destined for replacement (Manzano-Marín et al., 2018). Because the genes for the key nutritional molecule carbamoyl phosphate only exist on *S. endoviburni* CPS, we expect that this endosymbiont is safe from extinction, at least for now.

## Materials and Methods

### Insect rearing

We used mealybugs from a colony reared to study the transmission of a selfish B chromosome (Vea et al., 2021). In brief, we initially obtained mealybugs from a glass house in the Royal Botanic Gardens of Edinburgh in Scotland; from these insects, we established a laboratory colony fed on sprouting potatoes at 25C on a 16hr light/8 hr dark cycle.

### Sequencing and assembly

Illumina and PacBio sequence reads were obtained from and processed as described in Vea et al., 2021. Illumina reads were quality trimmed using Trimmomatic v0.36 (minimum length=36 bp, sliding window=4 bp, minimum quality score=15 [ILLUMINACLIP:TruSeq3-PE:2:30:10 LEADING:3 TRAILING:3 SLIDINGWINDOW:4:15 MINLEN:36]) (Bolger et al., 2014). PacBio and Illumina reads were then assembled using Canu v1.6 (default parameters; Koren et al., 2017), resulting in 2,787 contigs and 440,161,839 bases. Contigs that appeared to be bacterial were extracted from the assembly using the SprayNPray software (Garber et al., 2022), and these putative endosymbiont contigs were then used to recruit Illumina and PacBio reads. Mapping of Illumina reads was carried out using Bowtie2 v2.3.4.1 (Langmead and Salzberg, 2012). Mapping of PacBio reads was done with BLASR v5.1 (Chaisson and Tesler, 2012). Using Unicycler v0.4.8 (default paramters; Wick et al., 2017), we then performed a hybrid assembly of the putative endosymbiont-affiliated PacBio and Illumina reads. Endosymbiont genomes have been made publicly available via FigShare: https://doi.org/10.6084/m9.figshare.24945384.v1.

### Phylogenomic analysis

Phylogenomic analyses were carried out using GToTree v1.5.38 (Lee, 2019). Phylogenomic tree construction was carried out in RaxML, with 100 bootstraps (-N 100), the PROTCAT model for amino acid substitution, and the BLOSUM 62 amino acid matrix (-m PROTCATBLOSUM62) (Stamatakis, 2014).

### Functional annotation and pseudogene identification

We annotated each endosymbiont genome using Prokka (Seeman, 2014), which also predicted genes and open reading frames using a variety of software, including Prodigal (Hyatt et al., 2010) and RNAmmer (Lagesen et al., 2007). Protein coding genes were also annotated using the GhostKOALA annotation server (Kanehisa et al., 2016). Pseudogenes were identified using the software Pseudofinder (Syberg-Olsen et al., 2020). Annotation data were consolidated with the pseudogene predictions and organized in biosynthetic pathways using a semi-automated approach, which included custom python scripts and visual inspection.

We identified putative bacteria-to-insect horizontal gene transfers (HGTs) using the SprayN-Pray software (Garber, 2022) combined with previous published genomes (Husnik et al., 2016). Briefly, SprayNPray identified eukaryotic contigs using a combination of metrics, including contig length, coding density, and GC-content. Open reading frames (ORFs) from eukaryotic contigs were then compared against NBIC’s non-redundant (nr) database of proteins using DIAMOND (Buchfink et al., 2015), and top 100 matches evaluated. ORFs that recruited mostly (>50%) bacterial homologs were flagged as potential HGTs.

---

**Supplemental Figure 1:**
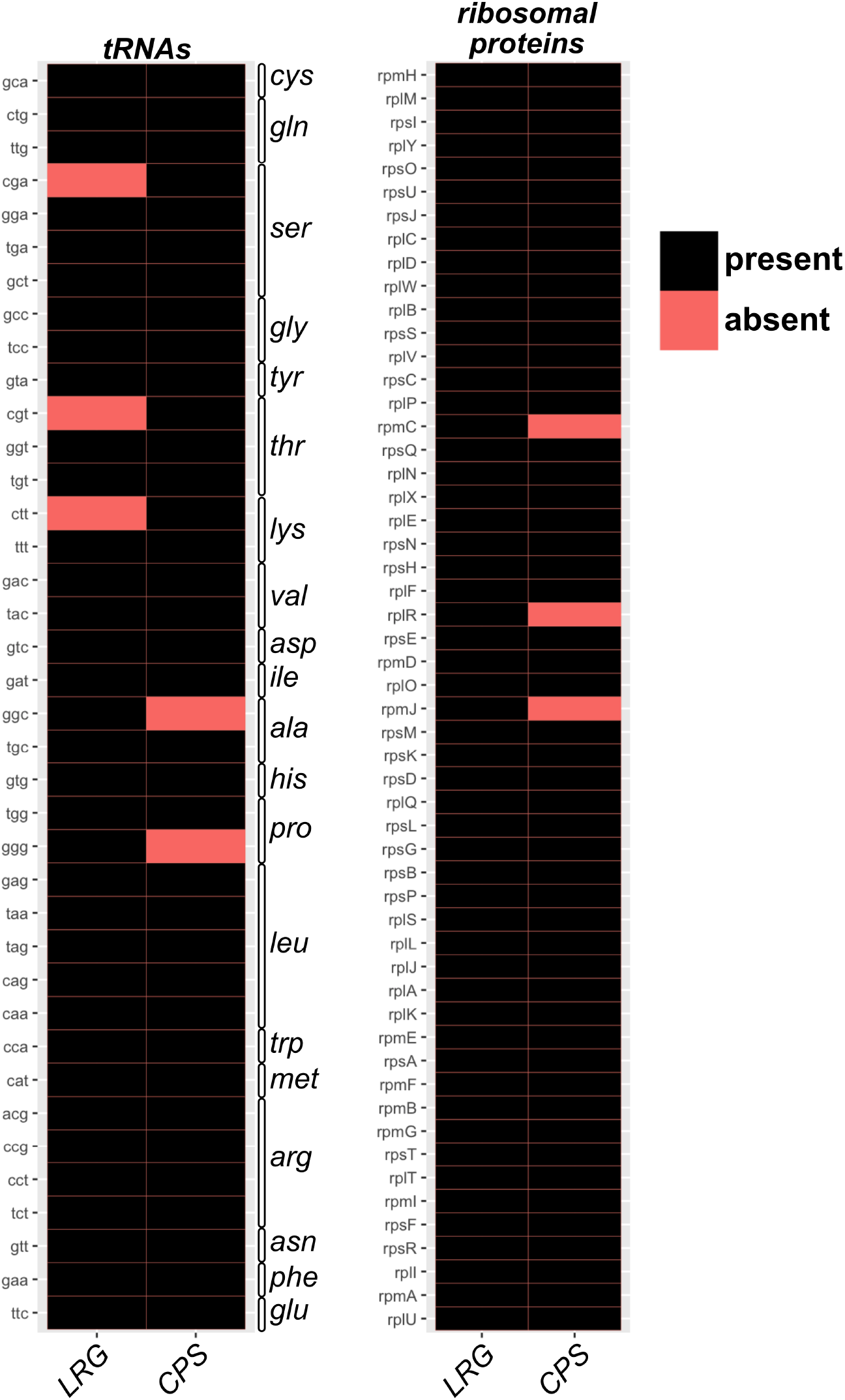
Presence/absence matrix showing the distribution of tRNAs and ribosomal proteins in the two Sodalis-related endosymbionts of P. viburni.

**Supplemental Figure 2:**
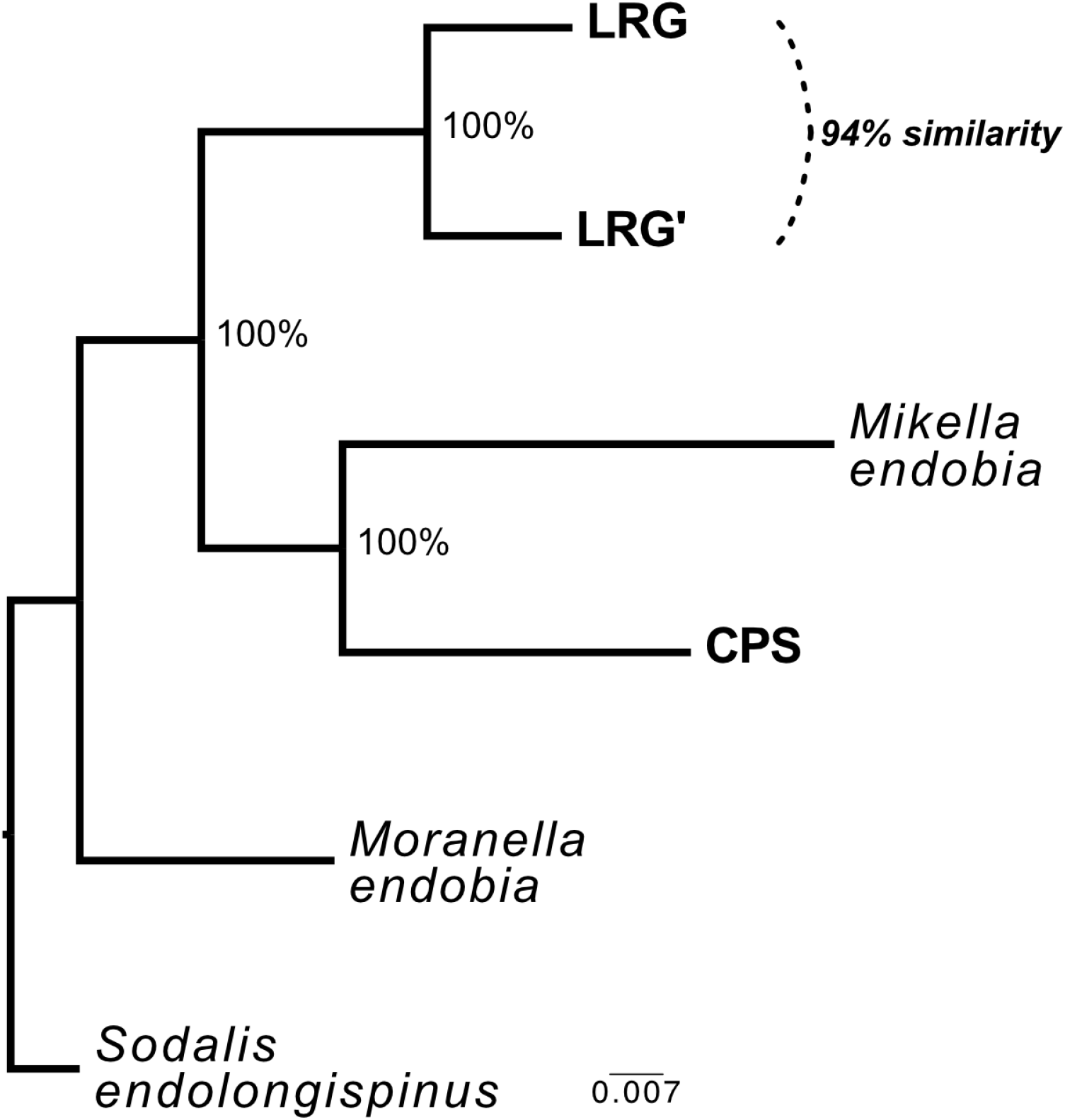
A phylogenetic tree based on a concatenated alignment of the 5S, 23S, and 16S rRNA genes. The two copies of the 16S gene encoded by S. endoviburni LRG are denoted by LRG and LRG’

## Notes

### Competing Interest Statement

The authors have declared no competing interest.

